# Identification of candidate genes for self-compatibility in perennial ryegrass (*Lolium perenne* L.)

**DOI:** 10.1101/2021.03.29.437398

**Authors:** Claudio Cropano, Chloé Manzanares, Steven Yates, Dario Copetti, Javier Do Canto, Thomas Lübberstedt, Michael Koch, Bruno Studer

**Affiliations:** Molecular Plant Breeding, Institute of Agricultural Sciences, ETH Zurich, Universitaetstrasse 2, 8092 Zurich, Switzerland; Deutsche Saatveredelung AG, Weissenburger Str. 5, 59557 Lippstadt, Germany; Department of Evolutionary Biology and Environmental Studies, University of Zurich, 8057 Zurich, Switzerland; Instituto Nacional de Investigación Agropecuaria (INIA), Ruta 5km 386, 45000 Tacuarembó, Uruguay; Department of Agronomy, Iowa State University, 716 Farm House Ln, 50011 Ames, USA

**Keywords:** self-incompatibility (SI), self-compatibility (SC), perennial ryegrass (*Lolium perenne* L.), quantitative trait locus (QTL) analysis, segregation distortion, fine mapping, candidate genes

## Abstract

Self-incompatibility (SI) is a genetic mechanism preventing self-pollination in approximately 40% of plant species. Two multiallelic loci, called *S* and *Z*, control the gametophytic SI system of the grass family (Poaceae), which contains all major forage grasses. Loci independent from *S* and *Z* have been reported to disrupt SI and lead to self-compatibility (SC). A locus causing SC in perennial ryegrass (*Lolium perenne* L.) was previously mapped on linkage group (LG) 5 in a F_2_ population segregating for SC. Using a subset of the same population (n=68), we first performed low-resolution quantitative trait locus (QTL) mapping to exclude the presence of additional, previously undetected contributors to SC. The previously reported QTL on LG 5 explained 38.4% of the phenotypic variation, and no significant contribution from other genomic regions was found. This was verified by the presence of significantly distorted markers in the region overlapping with the QTL. Second, we fine mapped the QTL to 0.26 cM using additional 2,056 plants and 23 novel sequence-based markers. Using an Italian ryegrass (*Lolium multiflorum* Lam.) genome assembly as reference, the markers flanking SC were estimated to span a ~3 Mb region encoding for 57 predicted genes. Among these, seven genes were proposed as relevant candidate genes based on their annotation and function described in previous studies. Our work is a step forward to identify SC genes in forage grasses and provides diagnostic markers for marker-assisted introgression of SC into elite germplasm.

**Key message:** A previously reported QTL conferring self-compatibility to perennial ryegrass was fine mapped to a 0.26 cM locus, corresponding to an estimated ~ 3 Mb physical region containing 57 predicted protein-encoding genes.

## Introduction

Several species of the grass family (Poaceae) are economically valuable forage crops and represent a fundamental component of our grasslands. Most forage grasses are obligate outcrossers due to a gametophytic self-incompatibility (SI) system that prevents inbreeding (Cornish et al. 1979). Unlike most self-incompatible species, SI in grasses is controlled by two multiallelic loci, called *S* and *Z*. Their existence is known since more than half a century (Hayman 1956; Lundqvist 1956) but, despite several mapping efforts (Voylokov et al. 1998; Thorogood and Armstead 2002; Bian et al. 2004; Hackauf and Wehling 2005; Kakeda et al. 2008; Shinozuka et al. 2010), the identity of the genes involved is unknown. To date, two genes encoding for proteins with a DUF247 domain on linkage group (LG) 1 and 2 are the most promising candidates for *S* and *Z*, respectively, in perennial ryegrass (*Lolium perenne* L.) (Shinozuka et al. 2010; Manzanares et al. 2016a).

There is a continued need to accelerate breeding in forage grasses for traits such as biomass yield and quality, seed yield, and disease resistance (Capstaff and Miller 2018). However, the genetic gain achieved in self-incompatible grasses with population breeding strategies has lagged behind compared to inbred crops such as maize (*Zea mays* L.), wheat (*Triticum aestivum* L.), and rice (*Oriza sativa* L.) (Laidig et al. 2014). This is partially caused by the inability to develop F_1_-hybrid cultivars and exploit the hybrid vigour that occurs when two genetically distant (and usually highly homozygous) parents are crossed. Thus, the possibility to overcome SI and exploit self-compatibility (SC) to transition forage grasses to an inbred line-based breeding system is gaining traction (Do Canto et al. 2016; Herridge et al. 2020). Inbreeding by self-pollination fixes favourable genetic variants and purges deleterious alleles with high efficiency, resulting in highly homozygous inbred lines (Jansky et al. 2016). Inbred lines belonging to different heterotic groups can be crossed to systematically assemble desirable combinations of genes and maximize heterosis.

Several routes to SC have been described in grasses: SC can arise from mutations at *S* and/or *Z* that disrupt the initial self/non-self recognition between pollen and stigma (Do Canto et al. 2016). Mutations in loci non-allelic to *S* and *Z* can also lead to SC, by interrupting the downstream cascade triggered by the initial self-recognition (Do Canto et al. 2016). Such putative mutations have been found in rye (*Secale cereale* L.) and perennial ryegrass (Voylokov et al. 1993; Egorova et al. 2000; Thorogood et al. 2005; Arias-Aguirre et al. 2013, Do Canto et al. 2018; Slatter et al. 2020). A detailed molecular understanding of how SC arises in SI grasses is lacking, but several studies have acknowledged that calcium (Ca^2+^) signaling plays an essential role in the recognition and/or inhibition of self-pollen (Yang et al. 2009; Chen et al. 2019). In fact, SI in grasses can be partially overcome by treating self-pollinated stigmas with chemical reagents affecting Ca^2+^ channeling across cell membranes (Klaas et al. 2011).

Two consecutive studies provided evidence on the location of a SC source segregating in a F_2_ population of perennial ryegrass. Using *in vitro* pollinations, Arias-Aguirre et al. (2013) observed a 1:1 segregation into two phenotypic SC classes: plants showing a 50% pollen compatibility where half of the self-pollen germinated and grew a pollen tube upon contact with the stigma, and 100% SC where all self-pollen showed a compatible reaction. It was concluded that SC was caused by a putative mutation in a major gametophytic gene mapped to a 16 cM locus on LG 5. Do Canto et al. (2018) fine mapped the position of the locus to a 1.6 cM region. Despite providing evidence of one major locus explaining the SC variation, these two studies did not exclude the presence of additional major effects on other LGs. In addition, the genetic region co-segregating with SC in Do Canto et al. (2018) was inferred based on genome sequence information from rice and Brachypodium (*Brachypodium distachyon* (L.) P. Beauv.) and still considerably large, making it difficult to identify putative candidate genes for functional validation.

Building on these previous studies, we aimed at providing additional evidence for the quantitative trait locus (QTL) on LG 5 being the sole cause for SC variation in this population. An additional objective was to increase the genetic mapping resolution achieved by Do Canto et al. (2018) substantially, to facilitate transitioning from a genetic to a physical map in ryegrass, and to select and prioritize candidate genes in the target region. A better understanding of the genetic causes of SC can help biologists to determine the pathways involved in the SI system of grasses. In addition, it paves the way for shifting the breeding of perennial grasses from population-based to pure line-based F_1_ hybrid strategies.

## Material and Methods

### Plant material

This study was entirely based on the perennial ryegrass F_2_ population segregating for SC described in Arias-Aguirre *et al.* (2013) and Do Canto *et al.* (2018). The population was obtained by self-pollinating a single F_1_ individual originating from the initial cross between a self-compatible genotype (selfed for five generations) and a self-incompatible individual of the VrnA mapping population (Jensen *et al.*, 2005). A subset of 74 individuals of the population used in Do Canto *et al.* (2018) was used as starting material for linkage map construction, low-resolution QTL mapping and the analysis of segregation distortion. A larger set of 2,056 individuals, in addition to the 1,248 reported in Do Canto *et al.* (2018), was used for the fine mapping.

### DNA extraction

All genomic DNA used in this study was extracted from powdered frozen leaf tissue using the Mag-Bind^®^ Plant DNA DS 96 Kit (Omega Bio-tek, Inc. Norcross, GA, USA) on a 96-well plate KingFisher Flex Purification System (Thermo Fischer Scientific, Waltham, MA, USA) following manufacturer’s recommendations. Genomic DNA was visualized on a 1% agarose gel and quantified with a NanoDrop 8000 spectrophotometer (Thermo Fisher Scientific, Waltham, MA, USA).

### QTL mapping and analysis of segregation distortion

#### Phenotyping for self-compatibility

The SC phenotypic data of the 74 F_2_ individuals was obtained from the *in vitro* pollination assays data reported in Do Canto *et al.* (2018). Briefly, the number of compatible pollen grains, usually translucent and with a bright pollen tubes growing towards the style, was counted on at least ten stigmas per plant. The data for each plant consisted of the overall mean percentage of self-compatible pollen, using the stigmas as subsamples. A chi-square goodness of fit test was performed to test if the phenotypic variation deviated from the hypothesis of SC being under the control of a single gametophytic gene. In that case, only heterozygote and homozygote plants for the SC allele (corresponding to the 50 and 100% phenotypic class) are expected in a 1:1 ratio in the F_2_.

#### Genotyping-by-sequencing, data processing and DNA variant calling

A genotyping-by-sequencing (GBS) library was prepared following the protocol reported in Begheyn *et al.* (2018) starting from genomic DNA of 74 individuals of the F_2_ population and sequenced using 125 bp single-end reads on an Illumina HiSeq 2500 platform at the Functional Genomics Center Zurich (FGCZ), Zurich, Switzerland. Sequencing reads were demultiplexed using Sabre (https://github.com/najoshi/sabre), allowing no mismatches. After demultiplexing, all adapters and barcodes were removed from the reads which were then trimmed to 100 bp. Variant calling was performed with the GBSmode pipeline (https://github.com/stevenandrewyates/GBSmode) using the Italian ryegrass (*Lolium multiflorum* Lam.) genome assembly as a reference (Copetti *et al.*, 2021).

#### Genetic linkage map construction

A genetic linkage map was constructed using the R package R/qtl (Broman *et al.*, 2003). Single nucleotide polymorphisms (SNP) resulting from the GBS variant calling were filtered for a 5% MAF, < 40% missing values, and distorted segregation from the expected 1:2:1 ratio in a F_2_ population. Individuals showing < 40% missing genotype calls were also filtered. In addition, pairs of individuals with more than 90% matching genotypes were identified and one was removed. As a result, 473 high-quality SNPs and 68 individuals were kept for the construction of the genetic map. Markers were grouped in LGs with the *formLinkageGroups* function with a maximum recombination rate of 0.35 and minimum −log_10_(*p*-value) (LOD) threshold of 6. The initial marker order and genetic distances were established using the Kosambi mapping function (*d* = (1/4) ln(1+2*r*/1–2*r*)), where *d* is the mapping distance and *r* is the recombination frequency (Kosambi, 1943). Singletons and double recombinations inflating the map size were identified and corrected using a graphical genotyping approach (Young and Tanksley, 1989). All singletons not followed by at least four markers of the same haplotype were turned into missing values and marker order and genetic distances were re-calculated. This process was repeated until all singletons and double recombinations within a window of four markers were removed. Linkage group numbers were assigned based on the physical position of their markers on the Italian ryegrass assembly (Copetti *et al.*, 2021).

#### QTL mapping and analysis of segregation distortion

The R package R/qtl was used to perform QTL mapping (Broman *et al.*, 2003). Missing values in the genotype table were filled with the *fill.geno* function and QTL mapping was carried out using the *mqmscan* function, which scans the genome with a multiple QTL model based on haplotype dominance and considers additive effects. After running 1,000 permutations with an assumed genotyping error rate of 0.05, a LOD of 2.81 was set as the QTL significance threshold. Segregation distortion was calculated using a chi-square goodness of fit test to identify markers that significantly differed from the 1:2:1 expected segregation. A Bonferroni-corrected significance threshold was set at a LOD of 3.97.

### Fine mapping

#### Marker development for fine mapping

To detect novel DNA sequence polymorphisms for marker development, 10 ng of genomic DNA from 50 individuals belonging to the collection of 2,056 F_2_ individuals were pooled and sequenced at low coverage in 2×150 bp mode on an S4 flow cell lane of an Illumina Novaseq instrument at FGCZ. The paired-end reads were mapped to an Italian ryegrass reference assembly (Copetti *et al.*, 2021) using bowtie2 (v3.5.1) (Langmead and Salzberg, 2012) and alignments were converted to BAM format with SAMtools (v1.10) (Li *et al.*, 2009). A BAM file containing reads that aligned on the scaffolds delimited by the markers flanking the SC locus in Do Canto *et al.* (2018) (G05_065 and G05_095) was imported into the Integrative Genomics Viewer software (Robinson *et al.* 2011) and polymorphisms were visually inspected. To develop PCR-based markers, a 50-120 bp of consensus sequence spanning a single polymorphism showing the expected 50% frequency was extracted and used as a template for primer design. Primers were designed with Primer-BLAST (Ye *et al.*, 2012) using 59-61 °C as optimal melting temperature (Tm). Primer sets with the lowest self- and self 3’-complementarity (self-binding affinity) were chosen and tested for specificity. Similarly, BAM files of 25 genotypes with a 50% SC phenotype and 25 genotypes with a 100% SC phenotype used for GBS were also used as a template for PCR-based marker development.

#### Large-scale genotyping based on High Resolution Melting Analysis

Genotyping was performed with High Resolution Melting Analysis (HRMA) on a LightCycler^®^ 480 System (Roche, Basel, Switzerland) using the same conditions as Do Canto *et al*, (2018). To identify additional individuals carrying a recombination, 2,056 F_2_ plants were genotyped with the markers G05_065 and G06_096. The recombinants were further genotyped with 23 newly validated polymorphisms (Supplementary Table 1). The genotypes of G05_065 and G06_096 and of the new markers were combined into a local linkage map using the OneMap software (Margarido *et al.*, 2007). Among the new markers, seven non co-segregating markers were used to obtain marker order and genetic distances by estimating the multipoint likelihood of all the possible orders using the *compare* function. The Kosambi mapping function was used for estimation of mapping distances. The markers within the SC locus were positioned by BLASTN on the Italian ryegrass genome assembly to obtain physical distances and extract information of the annotated protein-coding genes.

#### Phenotyping of the recombinants for self-compatibility

The individuals showing recombination between G05_065 and G06_096 were phenotyped using *in vitro* pollinations. Eight to ten mature virgin pistils were dissected from florets between 08:00 a.m. and 09:00 a.m. and placed on a petri dish containing a medium consisting of 2% agarose, 10% sucrose, and 100 ppm boric acid. From the same plants, four to five inflorescences with flowering florets were enclosed in paper bags prior to anthesis to collect pollen. Around noon, the paper bags were shaken and the pollen that fell at the bottom of the bag was sprinkled on the stigmas in the petri dishes. After a minimum of three hours, the pollinated stigmas were detached from the ovary with a razor blade, moved to a microscope slide, and submerged in a few drops of a staining solution containing 0.2% aniline blue and 2% K_3_PO_4_ (Martin, 1959). After covering with a cover slide, the samples were allowed to stain overnight. Pollen growth was detected using a Leica DM12000M UV light microscope (Leica Microsystems, Wetzlar, Germany). Plants were classified as either of the two SC phenotype classes segregating in the population (100% or 50% SC) according to the proportion of pollen grains showing a compatible reaction (Figure 1). In 100% SC plants, all pollen grains are translucent and displayed a clear pollen tube growing towards the style. In the 50% SC plants, nearly half of the pollen grains showed a short, thickened and bright blue pollen tube, with arrested growth upon contact with the stigma.

**Fig 1.**
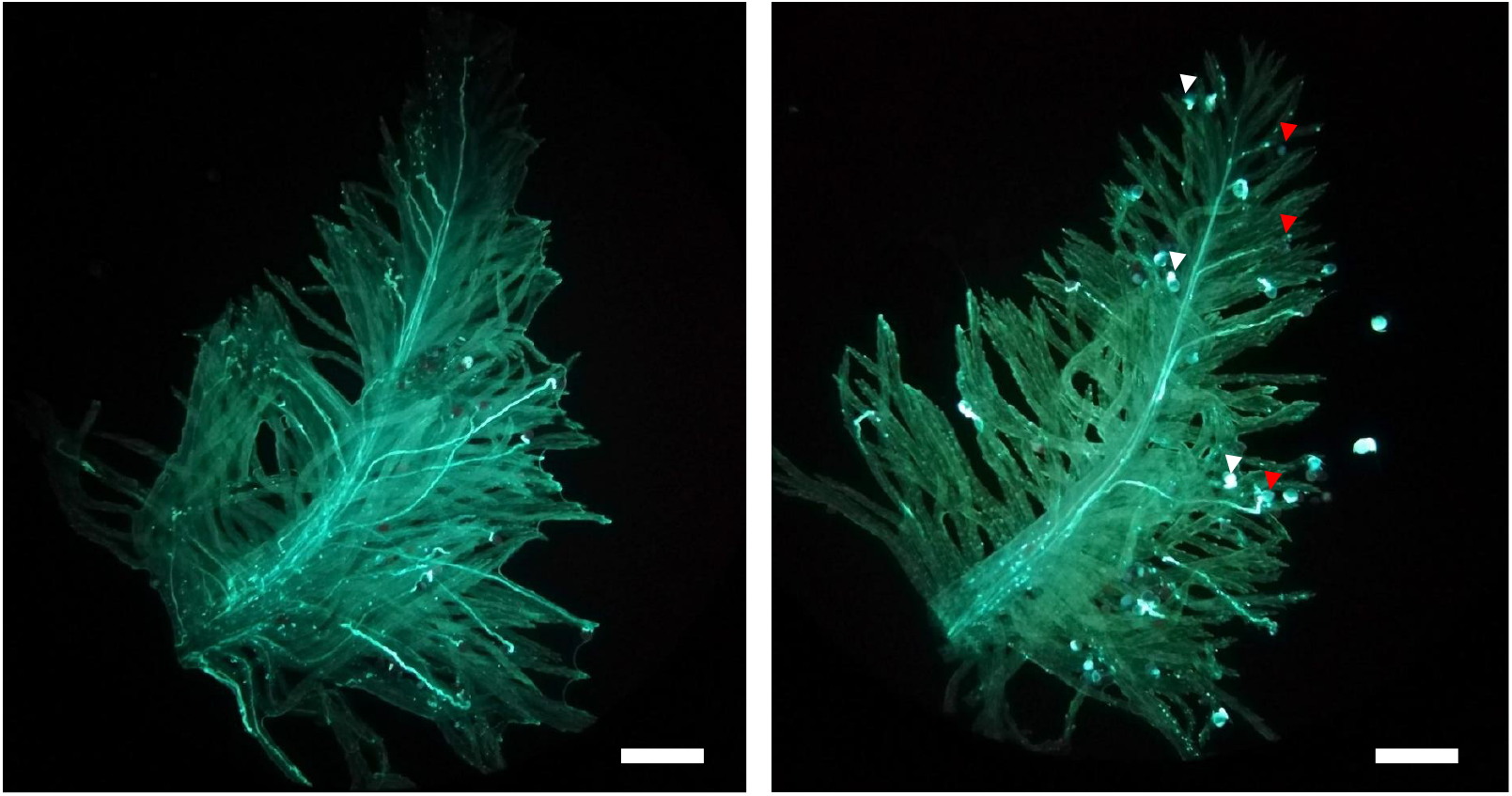
The two self-compatibility (SC) phenotypic classes segregating in the F_2_ population as visible in an *in vitro* pollination assay. a) In genotypes showing 100% SC, all pollen grains form a pollen tube that is able to elongate within the stigma reaching the ovary. b) In genotypes showing 50% SC, only half of the pollen grains penetrate the stigma and elongate a pollen tube (indicated by the red arrows). For the remaining half, pollen tube growth is aborted upon contact with the stigma and followed by a rapid deposition of callose at the pollen tube tip (white arrows). Bars: 400 μm.

### Identification of candidate genes in the genomic region co-segregating with self-compatibility

MCScanX (Wang *et al.*, 2012) was used to detect collinearity of the genes extracted from Italian ryegrass predicted in the genomic region co-segregating with SC with perennial ryegrass (Byrne *et al.*, 2015), barley (*Hordeum vulgare* L.) (Mascher *et al.*, 2017), rice (Ouyang *et al.*, 2007), and *Arabidopsis thaliana* (The Arabidopsis Genome Initiative, 2000) genome references by aligning their proteomes with BLASTP (Camacho *et al.*, 2009) at default parameters. BLASTP hits were retained, if they had an e value lower than 1e^-5^ and if they showed at least 80% and 60% (with perennial ryegrass, rice and barley, respectively) similarity to an Italian ryegrass gene. For *Arabidopsis thaliana*, the gene with highest similarity was chosen. Candidate genes were prioritized primarily based on functional annotation. Additional insights on their involvement in SC/SI were based on organ-specific gene expression data available on Genevestigator^®^ (Zimmermann *et al.*, 2005) for *Arabidopsis thaliana*, barley and rice orthologs. The pollen and stigma transcriptome data reported in Byrne *et al.* (2015) was used to assess if the perennial ryegrass genes present in the SC locus were differentially expressed during compatible/incompatible reactions.

## Results

### QTL mapping

#### Phenotypic segregation

Analysis of the *in vitro* pollination data for the 68 individuals from Do Canto et al. (2018) grouped the plants in two clear phenotypic classes (Figure 2a). The first class consisted of 43 plants with a mean compatibility of 90.5 (SD 11). Thirty plants were assigned to the second phenotypic class with a compatibility mean of 49% (SD 11.6). A chisquare test of goodness of fit showed no significant difference between observed and expected phenotypes of a monogenic trait under gametophytic control segregating in a 1:1 ratio in a F_2_ generation (*p*=0.128).

**Fig 2.**
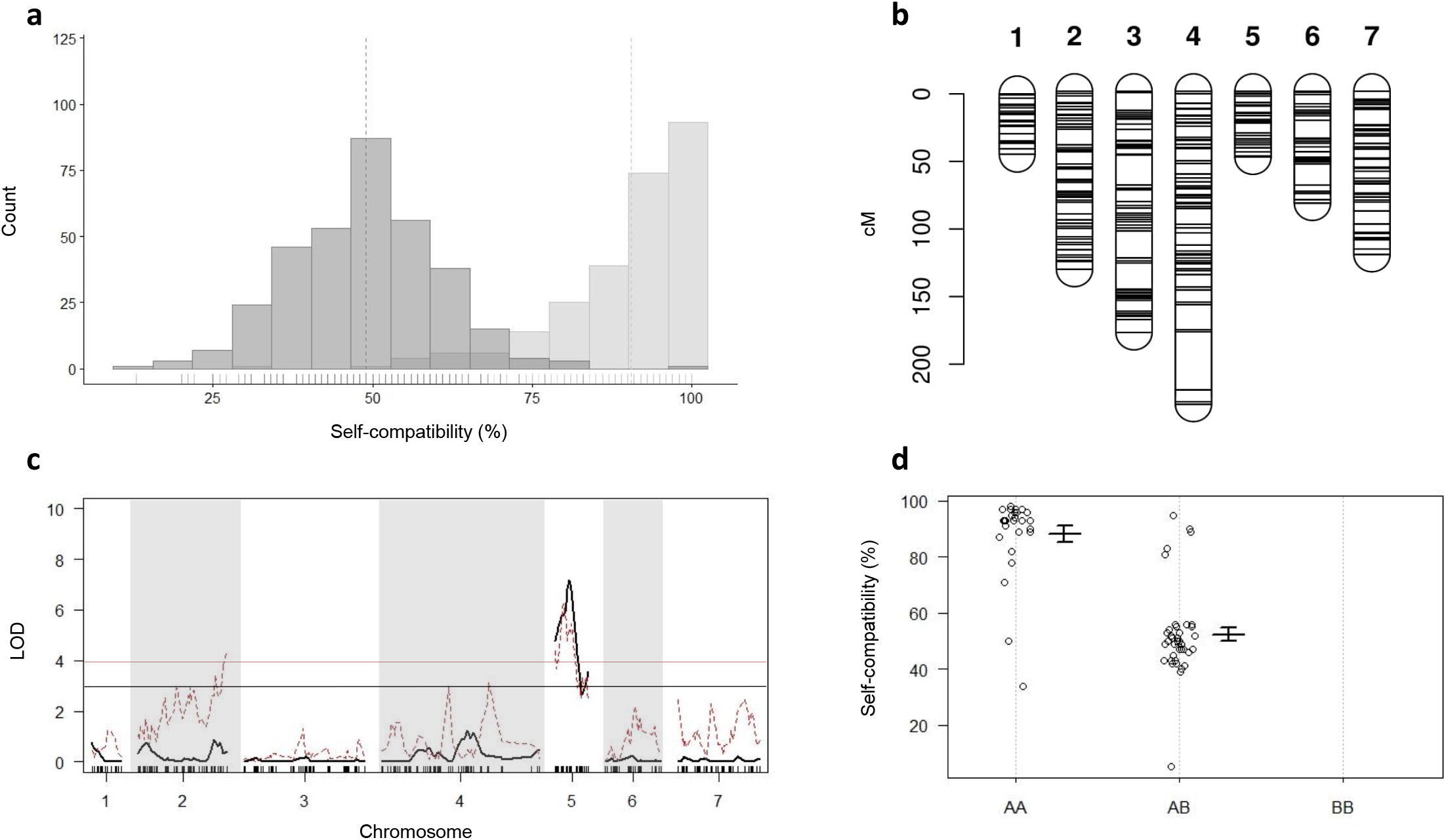
QTL analysis for self-compatibility (SC). a) The bi-modal phenotypic segregation for the SC phenotype of the 68 individuals used for the generation of the linkage map, QTL analysis and segregation distortion analysis. Data distributions and means (indicated by the vertical dotted lines) are displayed in dark grey for individuals with a 50% SC phenotype and light grey for the 100% self-compatible individuals. Small solid vertical lines above the horizontal axis indicate single data points. b) Genetic linkage map displaying the distribution of the 473 markers across the seven linkage groups of perennial ryegrass used for the QTL analysis. c) QTL analysis for SC (black line) and analysis of segregation distortion (red dotted line). Significance thresholds are indicated by the black and red horizontal lines for the QTL analysis and segregation distortion analysis, respectively. d) Marker effect plot for the most significant marker. AA represents the homozygous genotype for the SC phenotype, while AB and BB represent the heterozygous and homozygous genotypes for the self-incompatible phenotype, respectively. Plots show mean and standard deviation for the SC phenotype.

#### Genetic linkage and QTL mapping

Out of 10,487 SNPs resulting from the GBS variant calling, 473 passed the stringent quality filtering and were used for linkage map construction and QTL mapping. The distribution of the markers across the seven LGs ranged from 28 to 107, with an average of one marker every 1.8 cM and a total map length of 860.1 cM (Figure 2b, Table 1). The SC locus mapped to a single QTL on LG 5 with a maximum LOD value of 7.165 explaining 38.4% of the phenotypic variance (Figure 2c). An effect plot for the most significant marker, m1042, determined the contribution of the allelic states (A, H, B) to the respective phenotypes (Figure 2d). In the homozygous state for the allele contributed by the SC parent (AA), m1042 was associated with an increase of > 40% in self-compatible pollen in an *in vitro* self-pollination assay. The QTL found coincided with the QTL reported in Do Canto *et al.* (2018) as confirmed by the close physical proximity of m1042 and the two flanking markers, reported by Do Canto *et al.* (2018), on the Italian ryegrass reference assembly (data not shown).

**Table 1:** Descriptive statistics of the genotyping-by-sequencing-based genetic linkage map developed from 68 F_2_ individuals of perennial ryegrass segregating for self-compatibility. Length, average and maximum spacing for each linkage group are indicated in centimorgan (cM).

| Linkage group | Number of markers | Length (cM) | Average spacing (cM) | Maximum spacing (cM) |
| --- | --- | --- | --- | --- |
| **1** | 28 | 44.7 | 1.7 | 5.3 |
| **2** | 89 | 135.3 | 1.5 | 11.8 |
| **3** | 79 | 183.3 | 2.3 | 22.7 |
| **4** | 107 | 237.8 | 2.2 | 44.2 |
| **5** | 43 | 50 | 1.2 | 7.5 |
| **6** | 52 | 85 | 1.7 | 15.9 |
| **7** | 75 | 124 | 1.7 | 11.5 |
| **Overall** | **473** | **860.1** | **1.8** | **44.2** |

### Segregation distortion analysis

Analysis of segregation distortion performed using the genetic map employed for QTL mapping identified a region on LG 5 deviating from the 1:2:1 marker segregation expected in a F_2_ population. The region showed a 1:1 segregation in two genotypic classes (AA and AB) and overlapped with the region harboring the significant QTL for SC (Figure 2c). Similarly, four co-segregating markers (m1609, m113, m1780, m1440) at the end of LG 2 showed a significantly distorted segregation.

### Fine mapping and identification of candidate genes

To further dissect the SC locus, a total of 2,056 F_2_ plants were genotyped with the two markers G05_065 and G06_096 known to flank the locus in Do Canto *et al.* (2018). Thirty plants with recombination events between G05_065 and G06_096 were identified, nine of which survived and were further genotyped with the 23 newly developed markers (Supplementary Table 2). These additional genotyping data were used to generate a local linkage map and to estimate the genetic distances between those markers (Figure 3). The locus delimited by G05_065 and G06_096 spanned 1.44 cM, comparable to the genetic distance (1.6 cM) found by Do Canto *et al.* (2018).

**Fig 3.**
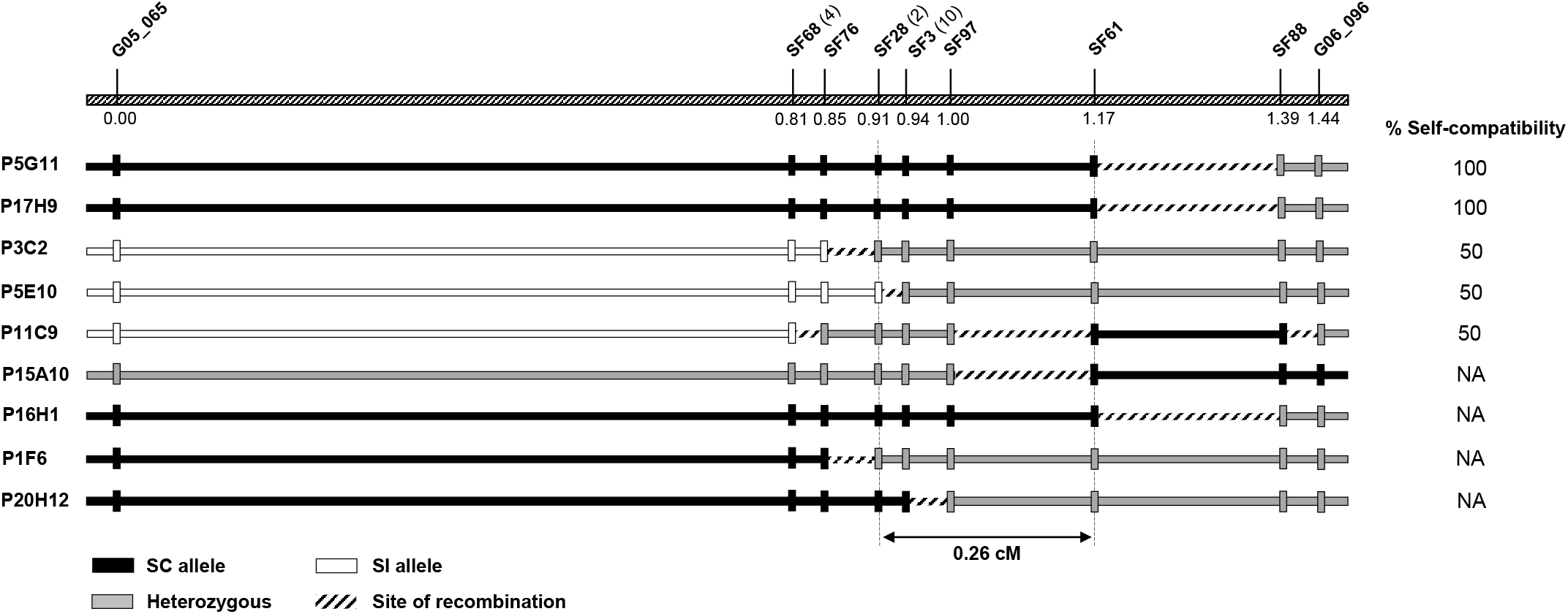
Fine mapping of the self-compatibility (SC) locus on linkage group 5. The top bar shows the genetic order and distances in centimorgan (cM) of the nine informative markers used for the calculation of genetic distances. For SF68, SF28 and SF3, the number of co-segregating markers is indicated in brackets. Genotype at each marker (vertical bars) and haplotype configuration (horizontal bars) for the nine individuals screened with G05_065 and G06_096 and the newly developed markers are shown. The column on the right-side reports the phenotypes, when available. The dashed vertical lines at SF28 and SF61 limit the SC locus.

Of the nine recombinants utilized for the calculation of genetic distances, five flowered and underwent *in vitro* pollinations, narrowing the region determining SC to 0.26 cM between markers SF28 and SF61. The position of the fine mapped locus was supported by a recombination in the interval between SF3 and SF28 found in P5E10 and by one individual with a recombination delimited by marker SF61 in P11C9 (Figure 3). Within the narrower region, one additional plant (P20H12) carried a recombination between SF3 and SF97, dividing the SC locus in two regions of 0.09 cM and 0.17 cM. However, P20H12 did not produce viable pollen, preventing phenotyping and the assignment of the SC co-segregating locus into one of the two sub-regions.

SF28 and SF61 mapped on two consecutive scaffolds of the Italian ryegrass genome assembly. This orthologous region spanned approximately 3 Mb and contained at least one sequence gap. In the interval, 57 protein-coding genes were predicted (Supplementary Table 3). The marker SF28 overlapped with *Lmu01_3856G0000030*, while SF61 mapped close to *Lmu01_702G0000540*, the most proximal gene inside the SC locus. The alignment of its proteome to the perennial ryegrass gene model revealed the presence of 27 highly similar genes distributed on 17 scaffolds. In barley, the region orthologous to the locus co-segregating with SC was approximately 3.6 Mb in size and contained 27 genes with *HORVU5Hr1G023880* and *HORVU5Hr1G024410* limiting the region on the SF28 and SF61 side, respectively. Fourteen genes had hits to the Italian ryegrass genes of the SC locus (Supplementary Table 3). The ortholog of the SC locus in rice was located between *LOC_Os12g36760* and *LOC_Os12g35670* on chromosome 12 and contained 76 genes, 16 of which had similarity to the genes predicted in Italian ryegrass. These orthologous regions from four different species allowed the compilation of a set of candidate genes based on their annotation, described function and information on organ-specific expression (Table 2).

**Table 2:**
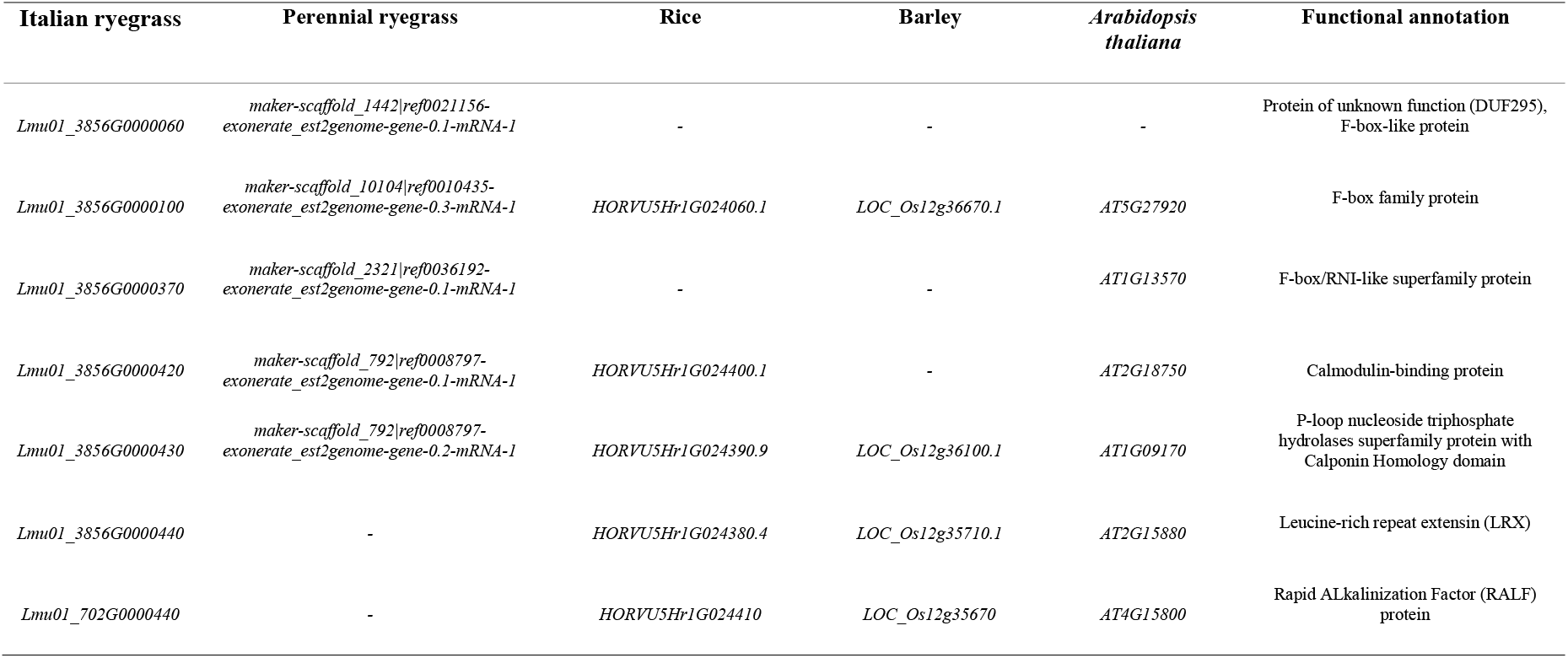
List of the seven prioritized candidate genes co-segregating with self-compatibility, their annotation and orthologs in perennial ryegrass (*Lolium perenne* L.), rice (*Oryza sativa* L.), barley (*Hordeum vulgare* L.) and *Arabidopsis thaliana*.

## Discussion

The identification of loci non-allelic to *S* and *Z* suggests that the grass SI system relies on a network of genes needed for the recognition, signal transmission and rejection or acceptance of self-pollen. How these loci interact with each other and their function in the SI downstream signaling cascade is yet to be determined. However, the SC locus studied here acts epistatically over *S* and *Z*: A pollen grain carrying the SC allele overcomes SI regardless of the composition at *S* and *Z*, and induces a 1:1 segregation in two SC classes: 50% and 100% SC (Arias-Aguirre *et al.*, 2013; Do Canto *et al.*, 2018). Genetically, this segregation pattern is only explained by the action of a single gametophytic gene. Under such a hypothesis, only the heterozygote and homozygote for the SC allele are transmitted to the F_2_ generation at a 1:1 ratio, resulting in the 50 and 100% pollen compatibility phenotypes. In our study, we observed the 1:1 phenotypic segregation pattern and, surveying a newly developed genome-wide linkage map, we found that the single QTL on LG 5 was not interacting with other genomic regions, confirming the single-locus inheritance.

Additional evidence was provided by the presence of a major region with distorted segregation in correspondence of the QTL on LG 5. In plants with a functional SI system, the segregation of a SC gene in certain crosses can lead to segregation distortion (Thorogood *et al.*, 2005). In our population, male gametes in the F_1_ plants carrying the non-SC allele are not transmitted to the F_2_ generation, causing a distorted segregation of markers linked to the SC causal gene. This has also been demonstrated recently by Slatter *et al.* (2020), where the QTL underlying the SC locus on LG 6 in perennial ryegrass overlapped with maximum marker segregation distortion. This has relevant implications for future studies aimed at discovering novel SC sources, as the identification of distorted regions can be used as a strategy to map their position in the genome. Mapping on the basis of segregation distortion represents a valid alternative to the low-throughput phenotyping provided by *in vitro* pollinations assays and it is not dependent on the duration of the flowering period. In addition, as no phenotyping is needed, it allows the screening of larger populations resulting in higher mapping resolution. A less relevant, yet significantly distorted region was also present on LG 2. There are several described biological reasons underlying marker segregation distortion (Xu *et al.*, 1997), thus it is difficult to identify its cause in this case.

After low-resolution QTL mapping, fine mapping resulted in the allocation of the locus to a 0.26 cM region between the markers SF28 and SF61, significantly reducing the extent of the previously described locus (Do Canto *et al.*, 2018) to a sub-cM region spanning ~ 3 Mb on the Italian ryegrass reference assembly. Importantly, there were nine additional markers that co-segregated with SF28, SF97, and SF61 due to lack of recombination events in the nine individuals that we identified. Such markers will be a valuable resource to further dissect genetically the SC locus, should additional F_2_ individuals be screened.

In the absence of a contiguous reference genome for perennial ryegrass, we cannot precisely define the genes present in the interval co-segregating with SC. However, given the high genome synteny with Italian ryegrass, barley and rice, it is relevant to assess and compare the genes annotated in the orthologous interval of their reference genomes as a proxy to suggesting candidate genes. On the Italian ryegrass reference assembly, the interval contains 57 protein-coding genes and we foresee to prioritize some of them as candidates responsible for SC based on their predicted function. The region showed conservation compared to other grasses, spanning a similar physical size on barley chromosome 5 and containing 27 genes. In rice, the SC locus ortholog was on chromosome 12. The wealth of functional information available for rice, barley, and *Arabidopsis thaliana* genes and similarity to other plant genes with known function allowed prioritization of genes for future identification of the SC determinant in perennial ryegrass.

Preliminary studies on the molecular determination of SI and SC suggested that Ca^2+^ signalling is involved in the recognition and/or inhibition of incompatible/compatible pollen. In fact, two independent expression studies registered an enrichment of transcripts during the SI response predicted to code for proteins containing calcium-binding domains in perennial ryegrass and sheepgrass (*Leymus chinensis* Trin.) (Yang *et al.*, 2009; Chen *et al.*, 2019). Two genes in the fine mapped SC locus region (*Lmu01_3856G0000420* and *Lmu01_3856G0000430*) encode for proteins with calcium-dependent functions. Their orthologs in *Arabidopsis thaliana*, *At2G18750* and *At1G09170*, encode for a calmodulin-binding protein and a kinesin motor protein with a calponin (calcium-binding protein) domain, respectively. No *At2G18750* expression in floral tissues has been described so far. However, *At1G09170* is co-expressed in mature pollen with several genes involved in mediating microtubule organization during pollen tube polar growth (Schneider and Persson, 2015). Interestingly, their orthologs in perennial ryegrass were reported to be up-regulated during compatible reactions (Byrne *et al.*, 2015), making them strong candidates for causing SC in our population.

Another relevant candidate gene is *Lmu01_3856G0000440*, whose function was annotated as a leucine-rich repeat extensin (LRX). Its ortholog in *Arabidopsis thaliana* (*At2G15880*) shows pollen-specific expression, and the rice ortholog (*LOC_Os12g356710*) shows high expression during pollen germination. Leucine-rich repeat extensins are cell wall proteins and part of a signalling system that transfers extracellular signals from the cell wall to the cytoplasm in order to regulate cell growth in vegetative tissue and in the pollen tube (Herger *et al.*, 2019). In several recent studies in *Arabidopsis thaliana*, *lrx* mutants show severe defects in pollen germination and pollen tube growth (Fabrice *et al.*, 2018; Sede *et al.*, 2018; Wang *et al.*, 2018). However, manipulating Ca^2+^ availability alleviates these defects, suggesting that LRX proteins can regulate Ca^2+^-related processes. In carrying out their regulatory function, LRX proteins interact with short cysteine-rich peptides known as RALFs (rapid alkanization factors). Interestingly, a gene encoding a RALF-like protein (*Lmu01_702G0000440*) is also present in the SC co-segregating region. RALFs induce a rapid alkalinization of extracellular space by increasing the cytoplasmic Ca^2+^ concentration that leads to calcium-dependent signalling events. They control several plant stress response mechanisms, growth, and development (Murphy and De Smet, 2014), and pollen-specific RALFs regulating pollen tube growth are described in tomato (*Solanum lycopersicum* L.) (Covey *et al.*, 2010) and *Arabidopsis thaliana* (Mecchia *et al.*, 2017; Moussu *et al.*, 2020). Both LRX and RALF-like-encoding genes are strong candidates for causing the inhibition of self-pollen tube. They could be a fundamental component of the process that inhibits self-pollen tube by transducing SI signals to activate the downstream pathway that modulates the Ca^2+^ gradient necessary for pollen tube growth. A disruptive mutation in such genes could interrupt the inhibitory mechanism, leading to SC.

Three other genes located in the narrower interval encode for F-box proteins (*Lmu01_3856G0000060*, *Lmu01_3856G0000100*, *Lmu01_3856G0000370*). *F-box* genes constitute one of the largest gene families in plants, involved in degradation of cellular proteins (Lechner *et al.*, 2006). F-box proteins form a subunit of the Skp1-cullin-F-box (SCF) complex that confers specificity for a target substrate to be degraded and are involved in important biological processes such as embryogenesis, floral development, plant growth and development, biotic and abiotic stress, hormonal responses and senescence, among others (Gupta *et al.*, 2015). *S*-locus *F-box* genes (*SFB* or *SLF*, depending on the family) are known as the male determinants of *S*-RNase based gametophytic SI systems (i.e. Rosaceae, Solanaceae, Scrophulariaceae, Rubiaceae). In these systems, the female determinant encodes for transmembrane *S*-RNase in the stigma with high cytotoxic activity for the pollen tube. In a non-self-pollination event, *SFBs* identify and degrade non-self *S*-RNases allowing fertilisation to occur unimpeded via the ubiquitin-26S proteasome pathway. Whereas in a self-pollination, self *S*-RNases evade degradation and exert cytotoxicity inside the pollen tube to inhibit its growth (Hua and Kao, 2008; McClure *et al.*, 2011). An *F-box* gene could also play a role in the complex downstream pathway that leads to rejection of self-pollen in grasses and a disruptive mutation might lead SC by halting its inhibiting function.

The genes discussed here are plausible candidates to cause SC, selected on the basis of their annotation and function described in previous studies. However, it is not possible to exclude the remaining genes in the genome region co-segregating with SC as potential candidates. Only additional, complementary analyses will allow the identification and confirmation of the molecular determinant of SC. Among these, targeted sequencing and sequence comparison of such genes in our SC population and diverse SI genotypes can contribute to reducing the number of candidate genes. Furthermore, techniques such as TILLING (Targeting Induced Local Lesions in Genomes) offer opportunities to screen for point mutations in candidate genes in populations mutagenized by chemical treatment and evaluate if those lead to a SC phenotype (Manzanares *et al.*, 2016b). Also, the recent advances in genome editing through CRISPR/Cas9 and its implementation in ryegrass (Zhang *et al.*, 2020) makes it an attractive alternative to achieve targeted gene inactivation of SC candidate genes to unequivocally determine their function.

## Conclusions

In this study, we provide additional evidence of a major QTL on LG 5 being solely responsible for SC in a perennial ryegrass population and demonstrate that segregation distortion can be an efficient strategy to identify loci conferring SC. Additionally, we developed new markers and fine mapped the previously described region from 1.6 cM down to 0.26 cM. Genome synteny to other grass species was used to determine a 3 Mb ortholog region in Italian ryegrass and to find the gene content in the region co-segregating with SC. From a total of 57 genes, seven encode for functionally relevant proteins and will be prioritized for validation as genetic determinants of SC. Furthermore, the genetic markers developed for fine mapping are tightly linked to the SC gene and can readily be converted into high throughput genotyping assays. They provide a convenient molecular tool to transfer SC to elite germplasm for the development of inbred lines-based breeding strategies.

## Acknowledgments

The authors wish to acknowledge Ingrid Stoffel-Studer and Dr. Timothy Sykes from the Molecular Plant Breeding group at ETH Zurich for assisting in DNA extraction and GBS library preparation, the Functional Genomic Center Zurich for sequencing service, the Zurich-Basel Plant Science Center for professional management of the PlantHUB programme and Nic Boerboom from DSV Zaden Nederland B.V for the helpful comments on the manuscript.

## Electronic supplementary material

### ESM_1.xlsx

Genotype and mapping data from 68 F2 individuals segregating for self-compatibility (SC), formatted as input for the R/qtl package. The first three rows indicate the marker names, their respective linkage group and genetic position in centimorgan. The column on the left indicates the averaged SC phenotypes from Do Canto et al., (2018) for each genotype.

### ESM_2.xlsx

Marker name, type of polymorphism, primer sequences and source of the markers used for High Resolution Melting Analysis.

### ESM_3.xlsx

High Resolution Melting Analysis of the markers G05_065 and G06_096 for the 2,056 F2 perennial ryegrass individuals screened for the identification of recombinants (A= homozygous for self-compatibility allele, B= homozygous for self-incompatibility allele, H=heterozygous)

### ESM_4.xlsx

High Resolution Melting Analysis of the markers used for fine mapping of the nine F2 individuals with at least one recombination event between G05_065 and G06_096 (A= homozygous for the self-compatibility allele, B=homozygous for the self-incompatibility allele, H=heterozygous).

### ESM_5.xlsx

Genes predicted in the Italian ryegrass orthologous region of the fine mapped self-compatibility locus and their orthologs in perennial ryegrass, barley, rice and *Arabidopsis thaliana* as resulting from BLASTP analysis.

## Declarations

### Availability of data and material

All data analyzed during this study are included in this published article and its supplementary information files. All raw sequencing data (deplexed GBS and pooled sequencing) are available at the Sequence Read Archive (https://www.ncbi.nlm.nih.gov/sra) under BioProject: PRJNA724991.

### Author contribution statement

Study conception: CC, CM, MK, TL and BS. Generation of plant material: JDC, TL. GBS library preparation, sequencing and bioinformatics: CC, CM and SY. Genetic map development, QTL mapping and analysis of segregation distortion: CC. Marker development, genotyping and marker data analysis: CC, CM, SY and DC. Comparative genomic analysis: CC and DC. Funding acquisition: MK, BS. Manuscript writing: CC, CM, DC and BS. All authors read and approved the final version of the manuscript.

